# Cinnamic acid and p-coumaric acid are metabolized to 4-hydroxybenzoic acid by *Yarrowia lipolytica*

**DOI:** 10.1101/2023.05.24.542070

**Authors:** Oliver Konzock, Marta Tous Mohedano, Irene Cibin, Yun Chen, Joakim Norbeck

## Abstract

The transition towards a bioeconomy requires the microbial production of various products from renewable resources such as lignocellulosic hydrolysate. *Yarrowia lipolytica* has been explored as a potential production host for flavonoid synthesis due to its high tolerance to aromatic acids and ability to supply malonyl-CoA. However, little is known about its ability to consume the precursors cinnamic and *p*-coumaric acid. In this study, we demonstrate that *Y. lipolytica* can consume these precursors through multiple pathways that are partially dependent on the cultivation medium. We constructed a collection of 15 P450 protein knock-out strains to identify the genes responsible for the reaction and identified YALI1_B28430g as the gene encoding for a protein with a trans-cinnamate 4-monooxygenase activity that converts cinnamic acid to *p*-coumaric acid and named it TCM1. *p*-Coumaric acid in turn is further converted to 4-hydroxybenzoic acid. Our findings provide new insight into the metabolic capabilities of *Y. lipolytica* and will be essential for the future construction of better flavonoid production strains.

## Introduction

Our current world economy is highly dependent on fossil fuels to produce a variety of products. Consequently, the excessive release of greenhouse gases is causing climate change and threatening the biodiversity of our planet. To move towards a more sustainable and environmentally friendly economy, microbial cell factories can play a key role (Vanholme et al. 2013).

Microbial cell factories are utilized for a constantly growing number of products, from plant-derived cancer medication (Courdavault et al. 2020) to bulk products like bioethanol (Robak and Balcerek 2020). To make the production processes economically and ecologically more feasible, lignocellulosic hydrolysates are used as a renewable alternative carbon and energy sources. However, those hydrolysates contain different microbial inhibitors e.g. phenolic compounds deriving from lignin such as cinnamic or *p*-coumaric acid (Palmqvist and Hahn-Hägerdal 2000). While a lot of research has been put into achieving tolerance to these inhibitors (Cunha et al. 2019), an even better approach would be to use them as precursors for valuable products.

Cinnamic and *p*-coumaric acid are the precursors for the flavonoids naringenin and pinocembrin, respectively. Both flavonoids have recently gained high interest for their antimicrobial, antioxidant, antitumor, and anti-inflammatory properties (Tungmunnithum et al. 2018). Naringenin and pinocembrin have both been produced in *Saccharomyces cerevisiae* (Koopman et al. 2012; Tous Mohedano et al. 2023) and *Escherichia coli* (Leonard et al. 2007). However, flavonoid biosynthesis requires malonyl-CoA as a precursor, a metabolite with a high flux during nitrogen starvation in the oleaginous yeast *Y. lipolytica*. Therefore, *Y. lipolytica* has already been explored as a production organism for different flavonoids e.g. naringenin (Wei et al. 2020) and liquiritigenin (Akram et al. 2021). Additionally, *Y. lipolytica* is known for its ability to consume a wide range of carbon sources e.g. different sugars, lipids, and alkanes (Ledesma-Amaro and Nicaud 2016). It is also highly tolerant to many of the inhibitors commonly found in hydrolysates (Konzock et al. 2021).

Most of the studies on flavonoid production have focused on increasing the production titers via a *push-and-pull* approach by testing different homologs of the pathway enzymes and adjusting their expression levels. However, very little research has been done to investigate any side reactions that might drain precursor supplies.

In this study, we investigated *Y. lipolytica*’s ability to consume the aromatic acids cinnamic and *p*-coumaric acid. We found that *Y. lipolytica* can metabolize cinnamic acid through at least two different pathways, that are furthermore medium-dependent. One pathway leads to the formation of *p*-coumaric acid and is catalyzed by the P450 protein encoded by YALI1_B28430g, herein named TCM1. *p*-Coumaric acid is then further converted to 4-hydroxybenzoic acid.

## Methods

### Strains and strain construction

The *Yarrowia lipolytica* strain OKYL029 is based on the W29 background strain (Y-63746 from the ARS Culture Collection, Peoria, USA; a.k.a. ATCC20460/ CBS7504) and was previously engineered to prevent filamentous growth by deletion of MHY1 (Konzock and Norbeck 2020) (MATa Δku70::Cas9::DsdA Δmhy1). Deletion strains were based on OKYL029 and constructed by removing the open reading frame of the corresponding gene using the EasyCloneYALI toolbox (Holkenbrink et al. 2018). Briefly, the target gene was targeted by two gRNAs to induce DNA double-strand breaks which were repaired by providing the cell with a 100 bp repair fragment for homologous recombination. Sequences of the gRNAs, repair fragments and validation primers can be found in the supplementary information.

Transformations were performed using a lithium acetate heat shock protocol according to EasyCloneYALI toolbox (Holkenbrink et al. 2018).

### Media

Unless differently stated, Delft media with a C/N ratio of 18 and LPU media with a C/N ratio of 200 was used.

Delft media (C/N 18) consisted of 7.5 g/L ammonium sulfate [Sigma Aldrich, 7783-20-2], 14.4 g/L potassium phosphate monobasic [Fisher Scientific, 7778-77-0], 0.5 g/L magnesium sulfate heptahydrate [Merck, 10034-99-8], 33 g/L D(+)-Glucose monohydrate [Avantor, VWR, 14431-43-7], 2 mL/L trace metals solution, 1 mL/L vitamin solution; pH adjusted to 5.5 with potassium hydroxide [Avantor, VWR, 1310-58-3] and is based on Verduyn et al. (1992).

The trace metal solution consisted of the following chemicals and their concentrations: 3 g/L Iron(II) sulfate heptahydrate (FeSO_4_*7 H_2_O) [Sigma Aldrich, 7782-63-0], 4.5 g/L Zinc sulfate heptahydrate (ZnSO_4_*7 H_2_O) [Sigma Aldrich, 7446-20-0], 4.5 g/L Calcium chloride dihydrate (CaCl_2_*2 H_2_O) [Sigma Aldrich, 10035-04-8], 1.0 g/L Manganese(II) chloride tetrahydrate (MnCl_2_*4 H_2_O) [Sigma Aldrich, 13446-34-9], 300 mg/L Cobalt(II) chloride hexahydrate (CoCl_2_*6 H_2_O) [Sigma Aldrich, 7791-13-1], 300 mg/L Copper(II) sulfate pentahydrate (CuSO_4_*5 H_2_O) [Sigma Aldrich, 7758-99-8], 400 mg/L Sodium molybdate dihydrate (Na_2_MoO4*2 H_2_O) [Sigma Aldrich, 10102-40-6, 1.0 g/L Boric acid (H_3_BO_3_) [Sigma Aldrich, 10043-35-3], 100 mg/L Potassium iodide (KI) [Sigma Aldrich, 7681-11-0], and 19 g/L Disodium ethylenediaminetetraacetate dihydrate (Na_2_EDTA*2 H_2_O) [Sigma Aldrich, 6381-92-6].

Vitamin solution consisted of 50 mg/L d-biotin [Sigma Aldrich, 58-85-5], 1.0 g/L D-pantothenic acid hemicalcium salt [Sigma Aldrich, 137-08-6], 1.0 g/L thiamin-HCl [Sigma Aldrich, 67-03-8], 1.0 g/L pyridoxin-HCl [Sigma Aldrich, 58-56-0], 1.0 g/L nicotinic acid [Sigma Aldrich, 59-67-6], 0.2 g/L 4-aminobenzoic acid [Sigma Aldrich, 150-13-0], 25 g/L myo-Inositol [Sigma Aldrich, 87-89-8].

Lipid production media (LPU) (C/N 200) consisted of 1.5 g/L yeast extract [Merck, 8013-01-2], 0.85 g/L, casamino acids [Formedium, CAS01], 1.7 g/L Yeast Nitrogen Base without amino acids and ammonium sulfate [Formedium, CYN0502], 5.1 g/L potassium hydrogen phthalate [Merck, 877-24-7] buffer adjusted to pH 5.5 with potassium hydroxide [Avantor, VWR, 1310-58-3], 110 g/L D(+)-Glucose monohydrate [Avantor, VWR, 14431-43-7], and 0.5 g/L urea [Sigma Aldrich, 57-13-6] (Tsakraklides et al. 2018).

Media with different nitrogen sources or C/N ratios only differ in the amount of nitrogen source and glucose added and are summarized in **Table 1**. Other media components were not changed. In the case of the addition of yeast extract and casamino acids to Delft media, the final concentration mimicked the concentration found in LPU media. The addition of vitamin and trace metal solution to LPU media mimicked the final concentration of Delft medium.

**Table 1:** Amounts of glucose and nitrogen source in different media to achieve different C/N ratios.

|  | Delft 3 AS | Delft 18 AS | Delft 116 AS | Delft 3 U | Delft 116 U | LP 3 U | LP 200 U |
| --- | --- | --- | --- | --- | --- | --- | --- |
| Urea [g/L] | 0 | 0 | 0 | 2.4 | 0.213 | 10.00 | 0.5 |
| Ammonium Sulfate [g/L] | 5.28 | 7.5 | 0.471 | 0 | 0 | 0 | 0 |
| Glucose [g/L] | 7.2 | 30 | 25 | 7.2 | 25 | 30 | 100 |

YPD plates contained 20 g/L peptone from meat [Merck, 91079-38-8], 10 g/L [Merck, 8013-01-2], 20 g/L D(+)-Glucose monohydrate [Avantor, VWR, 14431-43-7], and 20 g/L agar [VWR, 9002-18-0]. For selection, YPD plates were supplemented with 250 mg/L Nourseothricin [Jena Bioscience, 96736-11-7].

### Cultivation

Growth curves to determine the tolerance to organic acids were measured with the Growth Profiler 960 (Enzyscreen B.V., Heemstede, The Netherlands) using a standard sandwich cover with pins (CR1396b). Overnight cultures were inoculated from YPD plates into 5 ml of Delft media (without any organic acid added) in 50 mL falcon tubes and cultivated at 30°C and 200 rpm for 12 to 16 h. An Eppendorf tube with 800 μL of media was inoculated with the overnight culture to a starting OD_600_ of 0.05. The organic acids, trans-cinnamic acid [Sigma Aldrich, 140-10-3], trans-*p*-coumaric acid [TCI, 501-98-4], caffeic acid [Sigma Aldrich, 331-39-5], and trans-ferulic acid [Sigma Aldrich, 537-98-4] were added to the media from 25 mg/mL stock (in ethanol [Sigma Aldrich, 64-17-5]), and the final ethanol concentration was adjusted to 1.86% v/v for all conditions. The tube was vortexed and 150 μL were transferred to 4 wells of a 96-wells plate. Strains were cultivated at 30°C and 200 rpm and pictures to determine the OD_600_ were taken every 30 min.

Shake flask experiments to quantify the organic acid degradation were performed in 100 mL shake flasks with 10 mL media. Cultures were inoculated to a starting OD_600_ of 0.05 and incubated at 30°C and 200 rpm.

### HPLC quantification

The organic acid extraction and high-performance liquid chromatographer (HPLC) protocol is based on (Tous Mohedano et al. 2023). For the extraction of organic acids 0.5 mL of cell culture from the shake flask cultivation was mixed with 0.5 mL absolute ethanol (100%), vortexed for 5 min, and centrifuged at 4°C and 15000 xrf. The supernatant was transferred to glass vials and analyzed on an HPLC (Thermo Fisher Scientific) coupled to a photodiode array detector and equipped with a Discovery HS F5 150 mm x 46 mm column (particle size 5 μm; Sigma-Aldrich). Solvent A was 10 mM ammonium formate [Merck, 540-69-2] (pH 3, adjusted with formic acid [Sigma Aldrich, 64-18-6]) and solvent B was acetonitrile [Sigma Aldrich, 75-05-8]. The eluent flow rate was 1.5 mL/min. The elution gradient started with 5% solvent B (0–0.5 min), followed by a linear increase from 5% to 60% solvent B (0.5– 20.5 min), another linear increase from 60% to 100% solvent B (20.5–21.5 min), maintenance at 100% solvent B for 1 min (21.5–22.5 min), a linear decrease from 100% to 5% solvent B (22.5–23.5 min), and maintenance at 5% solvent B for 0.5 min (23.5–24 min). Cinnamic acid, ferulic acid, *p*-coumaric acid, and caffeic acid were detected at 289 nm, at retention times of 12.5, 9.75, 8.6, and 7.3 min, respectively (Figure S6). 4-hydroxybenzoic acid was detected at 6.3 min at 260 nm. Concentrations were calculated based on standard curves.

## Results

### Aromatic acid tolerance is partly based on degradation

In a previous study, we observed that *Y. lipolytica* was highly tolerant to the aromatic acid cinnamic acid (Konzock et al. 2021). We tested the growth tolerance of cinnamic acid and three additional aromatic acids, namely caffeic, ferulic, and *p*-coumaric acid. The four aromatic acids have a similar structure and only differ in the number and position of hydroxyl and methoxy groups. We observed a high tolerance of *Y. lipolytica* to all four acids up to a concentration of 2.4 mM (**Figure 1 A**).

**Figure 1:**
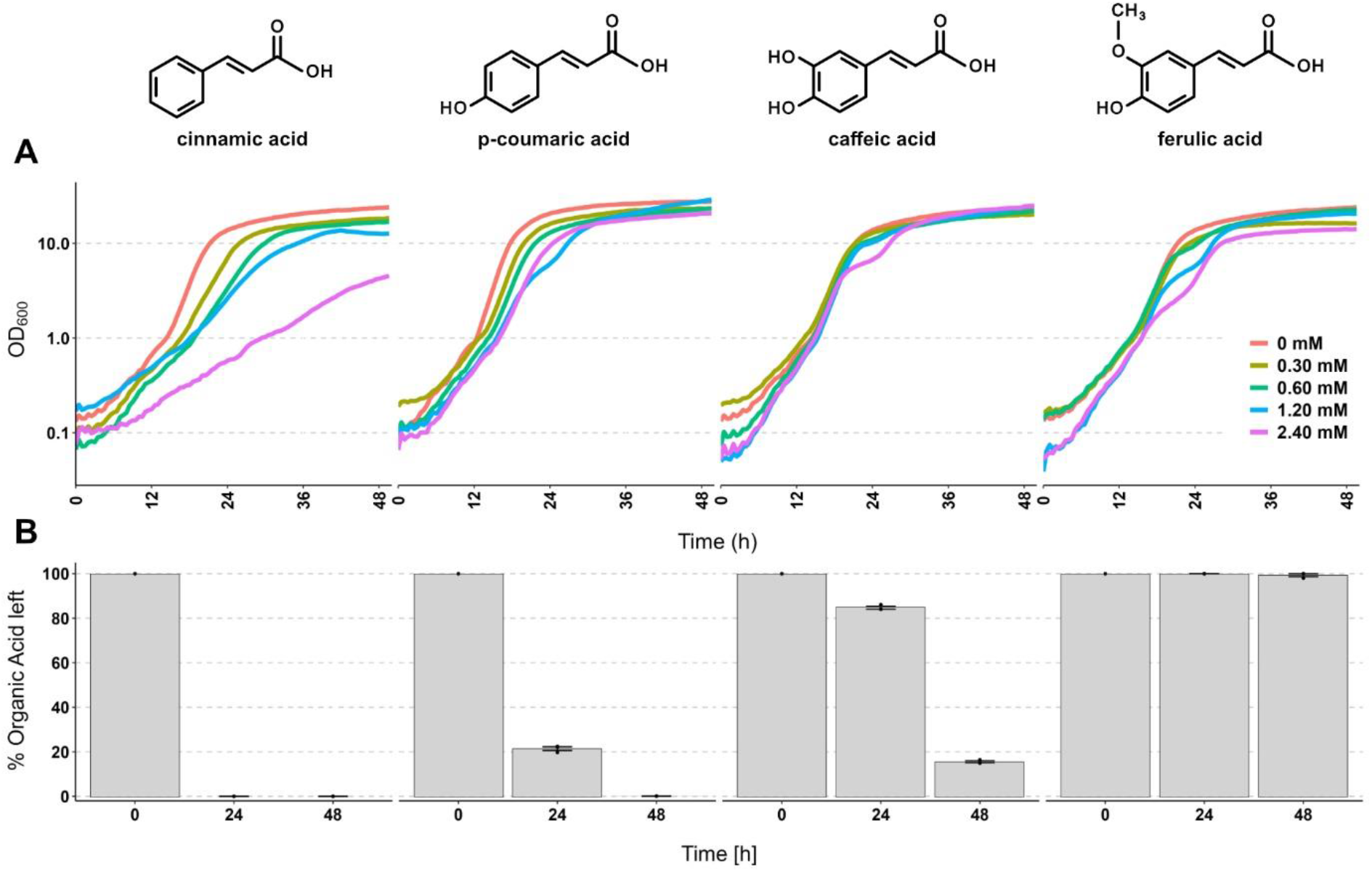
Tolerance to and degradation of different aromatic acids by *Y. lipolytica*. **A)** Growth over time of strain OKYL029 in Delft media with different starting concentrations of the aromatic acids cinnamic, *p*-coumaric, caffeic, and ferulic acid. Strains were cultivated in 96-well plates, and pictures to calculate the OD_600_ were taken every 30 min with the Growth Profiler 960. Lines represent the mean of four replicates (plot with standard deviations as ribbons in **Figure S2**). **B)** Degradation of the four acids after 24 and 48 hours. Strains were cultivated in 10 mL media in 100 mL shake flasks containing 125 mg/L of each acid (=0.843 mM cinnamic acid, 0.761 mM p-coumaric acid, 0.693 mM caffeic acid, 0.644 mM ferulic acid). Bars and error bars show the mean and standard deviation of the remaining organic acid (in %) of three replicates, respectively. The 0 h timepoint represents the organic acid concentration in the media before inoculation and was only measured once

Since *Y. lipolytica* is known to be able to consume a wide range of carbon sources (Ledesma-Amaro and Nicaud 2016), we measured the concentration of the four aromatic acids after 24 and 48 hours of cultivation (starting concentration 125 mg/L) (**Figure 1B**). We found that cinnamic, *p*-coumaric, and caffeic acid are all substantially degraded over time. After 24 h cinnamic acid is completely degraded, while 20% and 80% of *p*-coumaric and caffeic acid are remaining, respectively. After 48 h about 18% of caffeic acid remain and *p*-coumaric acid could not be detected anymore. We did not observe any degradation of ferulic acid during the two days of cultivation. We also tested the cinnamic acid degradation in a different strain isolate (A101.1.31, a UV-mutant from wild-type strain A101 (Wojtatowicz et al. 1991)) and confirmed a similar ability, indicating that is a conserved trait and not W29 strain-specific (**Figure S1**).

These results indicate that *Y. lipolytica*’s high tolerance to aromatic acids could be partly explained by its ability to degrade these compounds.

### *Cinnamic acid is degraded to* p*-coumaric acid and further degraded to 4-hydroxybenzoic acid*

Cinnamic and *p*-coumaric acid are the precursors for the flavonoids naringenin and pinocembrin, respectively. The ability to degrade these precursors is a significant drawback for *Y. lipolytica* as a host organism for flavonoid production. Therefore, we investigated the degradation pathway of cinnamic and *p*-coumaric acid further, to find a way to reduce their degradation.

We conducted a time course experiment, monitoring the growth (OD_600_) and concentration of cinnamic and *p*-coumaric acid over time. We used two different media, Delft media and lipid production media (LPU) media. Delft media is a defined media with ammonium sulphate as a nitrogen source with a moderate carbon/nitrogen ratio (C/N ratio = 18). LPU media uses urea as a nitrogen source, is buffered by potassium hydrogen phthalate buffer and has a high C/N ratio (=200) to induce lipid production. For both media, we observed the degradation of cinnamic and *p*-coumaric acid (**Figure 2 A**). Interestingly, in LPU media we observed the production of *p*-coumaric acid during cinnamic acid degradation. This indicates that cinnamic acid can be degraded to at least two distinct products, one being *p*-coumaric acid and one unknown compound (**Figure 2 B**). Based on the maximum *p*-coumaric acid concentration of 24 mg/L (after 12 h), we estimate that 20-25% of the cinnamic acid was degraded via this pathway.

**Figure 2:**
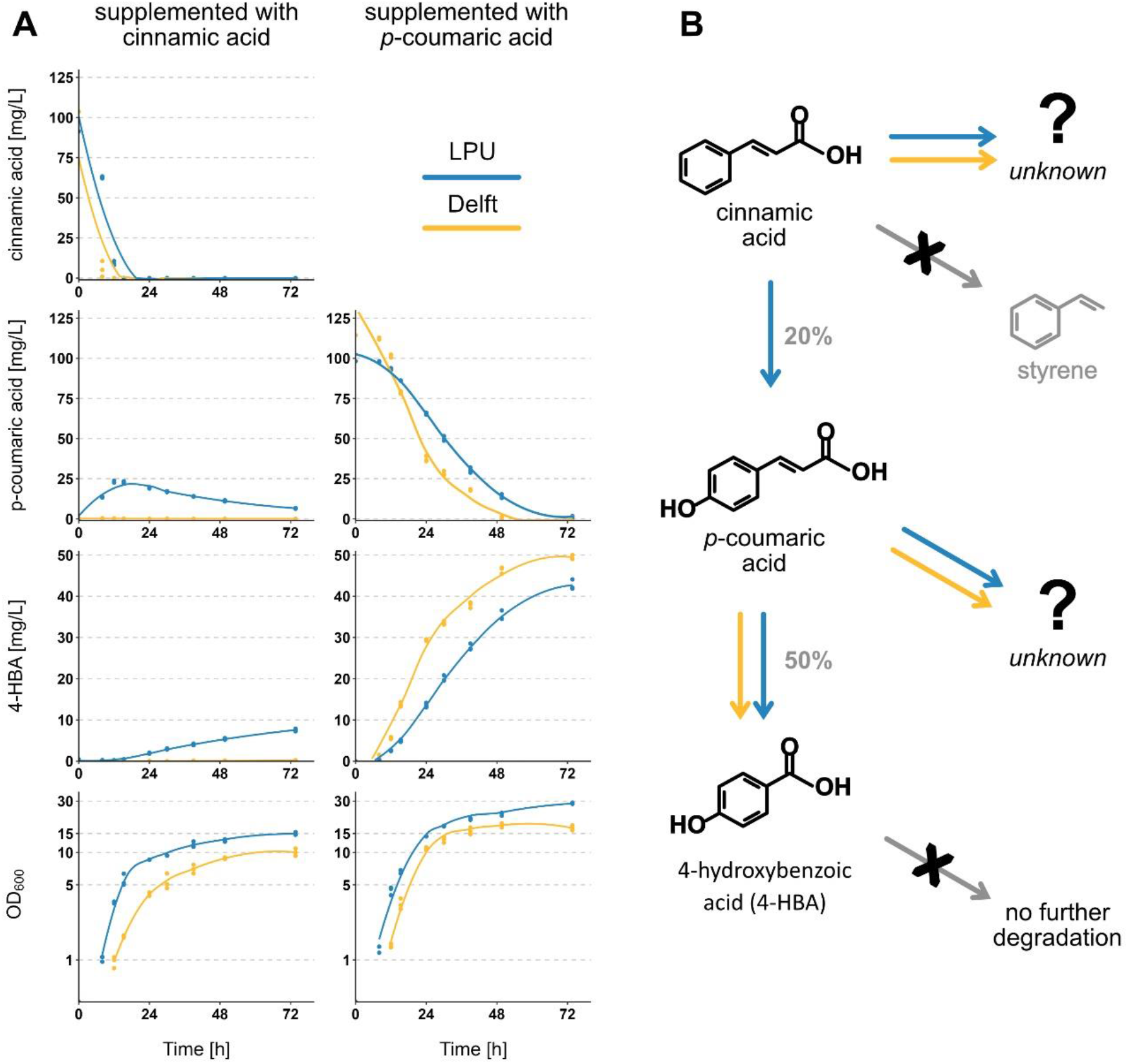
Cell growth and organic acid degradation over time. **A)** Strain OKYL029 was grown in 10 mL of either LPU or Delft media supplemented with 125 mg/L of cinnamic acid (left column) or *p*-coumaric acid (right column) in 100 mL shake flasks. Samples were taken for 74 h and analysed with the HPLC for cinnamic acid concentration (first row), *p*-coumaric acid (second row), 4-hydroxybenzoic acid (third row), and growth (last row). Dots are single data points; the line shows the tendency line calculated with geom_smooth with span=0.8 in ggplot2 (Wickham 2016). **B)** Summary of proposed degradation pathways of cinnamic and *p*-coumaric acid in *Y. lipolytica*. Cinnamic acid and *p*-coumaric acid are degraded to unknown compounds in both, LPU and Delft media. Cinnamic acid is converted to *p*-coumaric acid in LPU media, which is further converted to 4-hydroxybenzoic acid (4-HBA) in both media. 4-HBA is not further degraded and cinnamic acid is not converted to styrene (indicated by gray arrow with black x). % next to arrows represent the conversion of the educt to the product of the reaction.

To identify the unknown compound, we conducted a literature search for known degradation products from cinnamic acid. In *S. cerevisiae* cinnamic acid can be degraded to styrene by the Ferulic acid decarboxylase Fdc1 and Phenylacrylic acid decarboxylase Pad1 (Mukai et al. 2010; Richard et al. 2015). There are no reported homologs of these genes in *Y. lipolytica*. Additionally, we were able to detect a styrene standard with our detection method but did not observe the corresponding peak in any of our samples.

Since the conversion from cinnamic acid to *p*-coumaric acid only requires the addition of a hydroxyl group to the aromatic ring of cinnamic acid, we hypothesized that an additional hydroxyl group at *p*-coumaric acid could form caffeic acid, as observed in *Streptomyces* species (Nambudiri et al. 1972; Sachan et al. 2006). However, our HPLC analysis did not show a peak corresponding to caffeic acid in any of the samples supplemented with *p*-coumaric or cinnamic acid at any time point. However, as we have shown before, caffeic acid can be further degraded and would not necessarily be accumulated.

Another degradation product of *p*-coumaric acid in *Streptomyces* species is 4-hydroxybenzoic acid (4-HBA) (Sutherland et al. 1983; Sachan et al. 2006). We identified an accumulation of 4-HBA over time in our samples supplemented with *p*-coumaric acid and those supplemented with cinnamic acid in LPU media (**Figure 2**). Based on the quantification we estimate that 50% of the supplemented *p*-coumaric acid is converted to 4-hydroxybenzoic acid. Additionally, we supplemented a culture with 4-HBA (125 mg/L) and found that 4-HBA is not further converted within 5 days in either medium (**Figure S3**).

Based on these results, we hypothesized that cinnamic acid could be converted to benzoic acid by a similar reaction. However, we were not able to detect a benzoic acid standard with our HPLC setup.

### Cinnamic acid degradation pathway is depending on media composition

Based on the results from our previous experiment, we aimed to further understand the influence of the different media components on the degradation of both cinnamic and *p*-coumaric acid. We compared the different nitrogen sources (ammonium sulphate [AS] and urea), different carbon/nitrogen ratios (C/N ratio), and the addition of either yeast extract and casamino acids (YaC) to Delft media or vitamin and trace metal solution (VaM) to LPU media (details in method section). In all media, cinnamic acid was degraded but through different routes (**Table 2**). Interestingly, the addition of yeast extract and casamino acids to Delft media resulted in the production of *p*-coumaric acid and 4-HBA. Those two products were not detected in any of the other Delft-based media. This suggests that the degradation pathway from cinnamic acid to *p*-coumaric acid can be induced by the addition of complex media components.

**Table 2:** Influence of media composition on cinnamic acid degradation. Strain OKYL029 was cultured in 10 mL media supplemented with 125 mg/L cinnamic acid in 100 mL shake flasks at 30°C and 220 rpm for 22 h. Cinnamic acid, *p*-coumaric acid and 4-HBA show the average concentration in mg/L ± standard deviation of duplicates. - indicates no detection of the compound in either of the replicates. AS ammonium sulphate, YaC yeast extract and casamino acids, VaM vitamin and trace metal solution.

| media base | nitrogen source | C/N ratio | additional supplements | acid added | OD<sub>600</sub> | cinnamic acid [mg/L] | *p*-coumaric acid [mg/L] | 4-HBA [mg/L] |
| --- | --- | --- | --- | --- | --- | --- | --- | --- |
| Delft | AS | 3 | | cinnamic acid | 5.0 ± 0.0 | - | - | - |
| Delft | AS | 18 | | cinnamic acid | 5.1 ± 0.2 | - | - | - |
| Delft | AS | 18 | YaC | cinnamic acid | 8.5 ± 0.0 | - | 15.2 ± 0.5 | 9.3 ± 0.6 |
| Delft | AS | 116 | | cinnamic acid | 2.4 ± 0.1 | - | - | - |
| Delft | urea | 3 | | cinnamic acid | 6.0 ± 0.3 | - | - | - |
| Delft | urea | 116 | | cinnamic acid | 2.4 ± 0.1 | 7.3 ± 5.6 | - | - |
| LPU | urea | 3 | | cinnamic acid | 22.5 ± 0.4 | 37.8 ± 2.1 | 10.8 ± 0.6 | - |
| LPU | urea | 3 | VaM | cinnamic acid | 23.1 ± 1.2 | 9.7 ± 0.7 | 10.9 ± 0.4 | - |
| LPU | urea | 200 | | cinnamic acid | 13.7 ± 0.4 | - | 24.2 ± 0.3 | - |

Additionally, in LPU media with a lower C/N ratio, we found some cinnamic acid left in the media, while all of it was consumed in the high C/N ratio media. Simultaneously, the cell growth (OD_600_) was higher in a low C/N ratio, suggesting that the difference in cinnamic acid consumption is not a result of the number of cells but of a metabolic change in the cells.

For the degradation of *p*-coumaric acid, we did not find any link to the tested media components (**Table S1**).

### Gene YALI1_B28430g encodes for a protein with a trans-cinnamate 4-monooxygenase activity that converts cinnamic acid to p-coumaric acid

Next, we tried to identify the gene responsible for the degradation of cinnamic acid to *p*-coumaric acid. The reaction from cinnamic acid to *p*-coumaric acid requires a trans-cinnamate 4-monooxygenase (EC 1.14.14.91). A suitable and well-characterized protein facilitating this reaction is C4H of *Arabidopsis thaliana* (gene AT2G30490) (Mizutani et al. 1997). Using a BLAST-search the protein sequence of AtC4H was compared against all the predicted proteins from the genome of *Y. lipolytica* to find proteins that could catalyze the same reaction (details in the supplementary). We constructed single knock-out strains of 15 of 17 target genes and tested them for their ability to degrade cinnamic acid in both LPU and Delft media. The construction of knock-out strains of YALI1_C14106g and YALI1_B07094g was not successful, suggesting that they might be essential for growth. The growth of all other deletion strains was similar to the parental strain (**Figure S4**)

In Delft media, all knock-out strains continued to degrade cinnamic acid (**Figure S5**). Similarly, in LPU medium all strains consumed cinnamic acid (**Table 3**). However, one of the knock-out strains (ICYL09, ΔYALI1_B28430g) did not form any *p*-coumaric acid anymore (or 4-HBA). These results show, that YALI1_B28430g encodes for a protein with a trans-cinnamate 4-monooxygenase activity that converts cinnamic acid to *p*-coumaric acid. We hereby suggest naming this gene TCM1. It furthermore confirms our previous conclusion, that a second pathway for the degradation of cinnamic acid exists in *Y. lipolytica*.

**Table 3:** Degradation behaviour of different deletion strains. Shown are the strain name, the deleted gene name, and the protein name of the corresponding protein. Strains were cultured in 10 mL LPU media supplemented with 125 mg/L cinnamic acid in 100 mL shake flasks at 30°C and 220 rpm for 22 h. Cinnamic and p-coumaric acid show the average concentration in mg/L ± standard deviation of duplicates. 4-HBA was not detected in any of the samples.

| strain | deletion | protein name | OD<sub>600</sub> | cinnamic acid [mg/L] | *p*-coumaric acid [mg/L] |
| --- | --- | --- | --- | --- | --- |
| OKYL029 | none | | 13.7 ± 0.4 | 0.0 ± 0.0 | 24.2 ± 0.3 |
| ICYL06 | YALI1_A15544g | Alk7 | 8.9 ± 0.6 | 9.1 ± 0.4 | 14.1 ± 0.6 |
| ICYL07 | YALI1_F02132g | uncharacterized Protein | 9.1 ± 0.8 | 11.5 ± 2.9 | 13.7 ± 2.3 |
| ICYL08 | YALI1_F05415g | Alk2 | 8.4 ± 0.1 | 13.0 ± 0.0 | 12.3 ± 1.0 |
| ICYL09 | YALI1_B28430g | Cytochrome P450 | 8.4 ± 0.0 | 13.7 ± 0.5 | 0.0 ± 0.0 |
| ICYL10 | YALI1_A21175g | Cytochrome P450 | 10.2 ± 0.1 | 5.7 ± 0.7 | 16.0 ± 0.7 |
| ICYL11 | YALI1_B18347g | Alk5 | 8.8 ± 0.0 | 10.7 ± 1.6 | 13.4 ± 0.9 |
| ICYL12 | YALI1_E27649g | Alk3 | 10.5 ± 0.2 | 2.4 ± 2.4 | 22.6 ± 0.2 |
| ICYL13 | YALI1_E17508g | uncharacterized Protein | 10.1 ± 0.1 | 6.2 ± 0.0 | 16.4 ± 0.5 |
| ICYL14 | YALI1_B27057g | Cytochrome P450 | 9.5 ± 0.8 | 7.7 ± 2.8 | 16.6 ± 1.9 |
| ICYL16 | YALI1_B02833g | Alk6 | 10.2 ± 0.7 | 6.5 ± 0.9 | 14.1 ± 0.6 |
| ICYL17 | YALI1_A18344g | Cytochrome P450 | 8.4 ± 0.3 | 1.9 ± 0.6 | 15.3 ± 1.7 |
| ICYL18 | YALI1_E30815g | Cytochrome P450 | 8.4 ± 0.4 | 10.7 ± 1.1 | 12.8 ± 0.2 |
| ICYL19 | YALI1_C17031g | uncharacterized Protein | 8.1 ± 0.6 | 13.8 ± 2.1 | 8.2 ± 1.3 |
| ICYL20 | YALI1_B18311g | uncharacterized Protein | 8.3 ± 0.7 | 14.9 ± 1.5 | 11.5 ± 0.7 |
| ICYL21 | YALI1_B08321g | Cytochrome P450 | 10.1 ± 0.6 | 8.5 ± 1.2 | 12.1 ± 0.1 |

## Discussion

The growth experiments with different aromatic acids indicated that *Y. lipolytica* is highly tolerant toward these kinds of inhibitors as previously reported (Konzock et al. 2021). However, this previous study did not measure the aromatic acid concentrations over time and did not investigate possible degradation as the underlying cause of the high tolerance of *Y. lipolytica*. Our observation of aromatic acid degradation indicates that part of the tolerance could be based on this degradation. However, for ferulic acid we did not find degradation while simultaneously observing high tolerance, indicating that *Y. lipolytica* has indeed a high aromatic acid tolerance besides its ability to degrade some of these compounds.

We identified YALI1_B28430g as the gene encoding a protein with a trans-cinnamate 4-monooxygenase activity that converts cinnamic acid to *p*-coumaric acid. There are no specific publications linked to the gene on Uniprot exploring its function so far. Therefore, we suggest naming the gene YALI1_B28430g TCM1 according to its activity. Furthermore, our experiments with different growth media showed that TCM1-related enzyme activity is not observed in Delft media but that the addition of complex media components (yeast extract and casamino acids) can induce the TCM1-related enzyme activity.

The degradation of *p*-coumaric acid in *Y. lipolytica* has previously been observed in the context of resveratrol production which uses *p*-coumaric acid as a precursor (Sáez-Sáez et al. 2020). Although, the authors did not identify the degradation product they observed a reduced degradation of *p*-coumaric acid in YP medium containing complex nutrients. Our experiment with different media compositions indicates that the media composition can indeed affect the degradation of *p*-coumaric acid. However, we observed the opposite trend, with an additional degradation pathway of cinnamic acid to *p*-coumaric acid being induced by the addition of complex media compounds.

The identification of gene the gene TCM1 (YALI1_B28430g) to encode for the enzyme catalyzing the reaction from cinnamic to *p*-coumaric acid can be used for the development of microbial cell factories that use cinnamic acid as a precursor, e.g. pinocembrin or naringenin (Wei et al. 2020; Tous Mohedano et al. 2023). In the case of pinocembrin, a deletion of TCM1 would reduce the loss of precursor to the competing pathway, while overexpression of TCM1 could be beneficial for the production of naringenin, due to the formation of p-coumaric acid.

We have identified 4-hydroxybenzoic acid (4-HBA) as a degradation product of *p*-coumaric acid. The reaction can be catalyzed by a 4-coumarate-CoA ligase to form *p*-coumaroyl-CoA followed by the conversion by a 4-hydroxy benzoyl-CoA thioesterase to 4-HBA. Based on the initial concentration of *p*-coumaric acid and the resulting 4-HBA concentration we estimated that 50% of the *p*-coumaric acid is converted to 4-HBA. We conclude that at least one more degradation pathway to an unknown product exists. In *Streptomyces* species caffeic acid is an additional product of *p*-coumaric acid degradation (Sachan et al. 2006). While we did not observe any accumulation of caffeic acid in our experiments, we also found that caffeic acid itself is degraded when supplemented with the media. *p*-Coumaric acid may be partially converted to caffeic acid, which could in turn be further degraded.

The identification of 4-HBA as a product of *p*-coumaric acid in *Y. lipolytica* can be further explored as a sustainable production alternative. 4-HBA and its derivatives have a wide range of applications, e.g. as conservatives in cosmetics and pharmaceuticals (Lenzen et al. 2019) but also as a raw material of dyes (Kitade et al. 2018). Mainly, 4-HBA is produced via the Kolbe-Schmitt reaction which converts phenol and potassium hydroxide to potassium phenoxide, which further reacts with carbon dioxide to form 4-HBA (Lindsey and Jeskey 1957). However, since this reaction is very energy-consuming (high temperatures and pressure) more sustainable alternative production methods have been explored. For example, *Burkholderia glumae* has been engineered to produce a high concentration of 4-HBA from *p*-coumaric acid (Jung et al. 2016), while *Pseudomonas taiwanensis* (Lenzen et al. 2019) and *Pseudomonas putida* (Yu et al. 2016) have been engineered for *de novo* production of 4-HBA. Additionally, different yeasts have been explored to produce 4-HBA, e.g. *S. cerevisiae* (Averesch et al. 2017) and *Pichia pastoris* (Inokuma et al. 2023). Additionally, in *Y. lipolytica* 4-HBA has been produced and used as a precursor to producing arbutin, a cosmetic used for skin-lightening (Shang et al. 2020). However, all fungal and some of the bacterial examples rely on the heterologous expression of a chorismate pyruvate-lyase (ubiC) that converts chorismite (an intermediate of the shikimate pathway) to 4-HBA. The identification of an alternative native pathway in *Y. lipolytica* that uses p-coumaric acid can be further explored and used to exceed previous production titers.

In summary, we described the degradation pathways of cinnamic and *p*-coumaric acid in *Y. lipolytica*. We identified that multiple pathways exist for both acids, that *p*-coumaric acid is converted to 4-HBA and that the P450 protein encoded by YALI1_B28430g catalyzes the reaction of cinnamic acid to *p*-coumaric acid. These results will be essential for the development of future flavonoid production platform strains in *Yarrowia lipolytica*.

## Declarations

## List of abbreviations

4-HBA: 4-hydroxybenzoic acid
HPLC: high-performance liquid chromatography
LPU: Lipid production media
OD_600_: Optical density at 600 nm

## Ethics approval and consent to participate

Not applicable

## Consent for publication

Not applicable

## Availability of data and materials

All raw data as well as the calculated means and standard deviation plotted in the shown figures can be found in the supplementary.

## Authors’ contributions

Conceptualization: OK and MTM

Funding acquisition: JN and YC

Investigation: OK, MTM, and IC

Methodology: OK, MTM and IC.

Supervision: OK

Writing –original draft: OK

Writing –review & editing: OK, MTM, JN

## Acknowledgements

Strain A101.1.31 was kindly donated by Prof. W. Rymowicz and Prof. M. Robak from Wroclaw University of Environmental and Life Sciences.

