## Supplementary for "Cinnamic acid and p-coumaric acid are metabolized to 4-hydroxybenzoic acid by *Yarrowia lipolytica*"

### Content

**Supplementary figures**

- Figure S1: Cinnamic acid degradation of strain A101.1.31.
- Figure S2: Tolerance to different aromatic acids by *Y. lipolytica.*
- Figure S3: Long-term stability of 4-hydroxybenzoic acid in *Y. lipolytica* cultures.
- Figure S4: Growth of the deletion strains in Delft media.
- Figure S5: Cinnamic acid consumption of knock-out strains in Delft media.
- Figure S6: HPLC chromatogram of the four aromatic acids caffeic, *p*-coumaric, ferulic and cinnamic acid.

**Supplementary table**

- Table S1: Influence of media composition on *p*-coumaric acid degradation.

**Additional supplementary**

- Protein BLAST details

### Supplementary figures

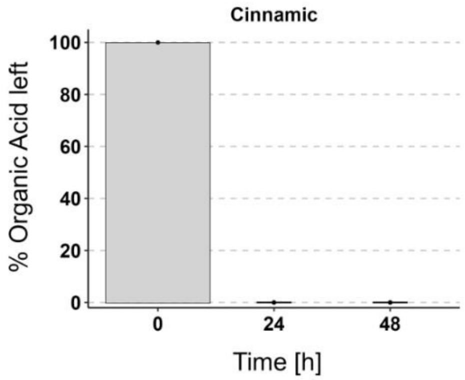


**Figure S1: Cinnamic acid degradation of strain A101.1.31.** The strain A101.1.31 (a UV-mutant from wild-type strain A101) was cultivated in duplicates in Delft media supplemented with 125 mg/L cinnamic acid at 30° and 220 rpm shaking. HPLC samples were taken after 24 and 48 h and analyzed. No Cinnamic acid peak was detected after 24 or 48 h. Other organic acids were not tested for this strain.

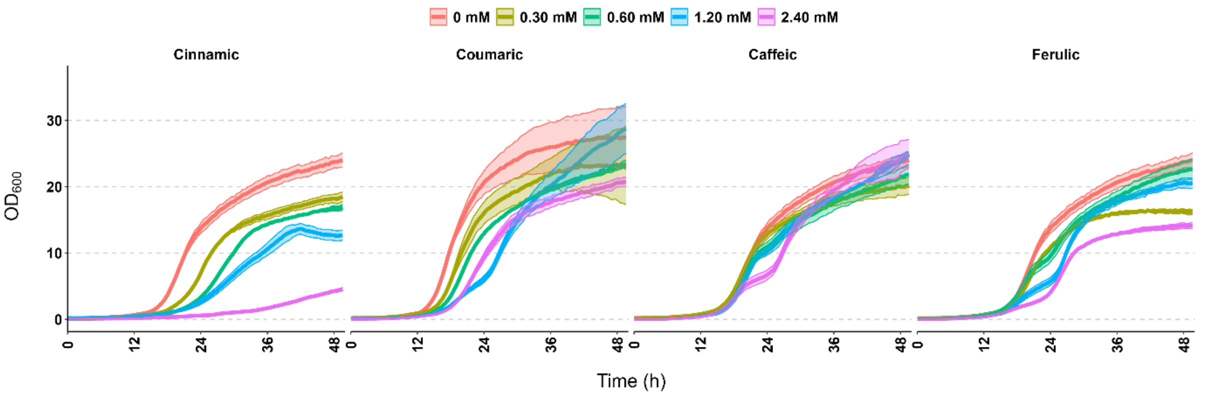


**Figure S2: Tolerance to different aromatic acids by *Y. lipolytica*.** Growth over time of strain OKYL029 in Delft media with different starting concentrations of the aromatic acids cinnamic, *p*-coumaric, caffeic, and ferulic acid. Strains were cultivated in 96-well plates, and pictures to calculate the $OD_{600}$ were taken every 30 min with the Growth Profiler 960. Lines and ribbons represent the mean and standard deviation of four replicates, respectively. The same data without ribbon is displayed in Figure 1A.

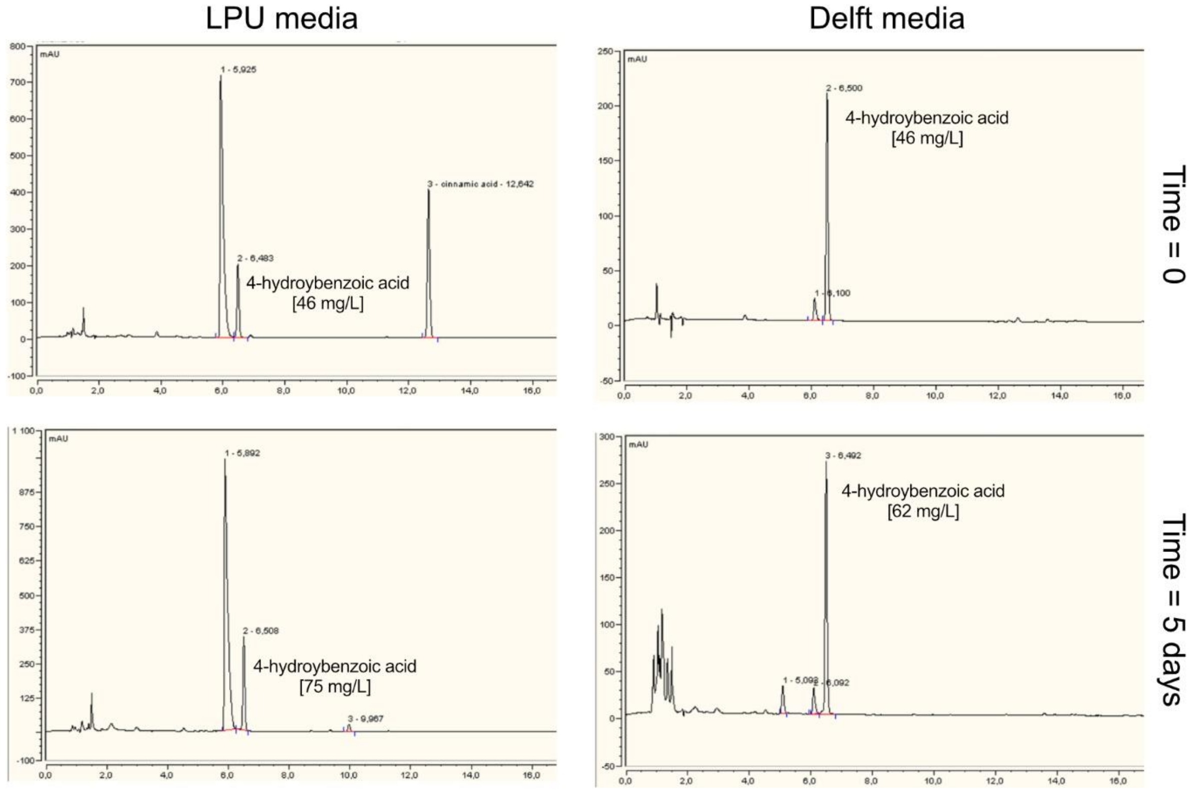


**Figure S3: Long-term stability of 4-hydroxybenzoic acid in *Y. lipolytica* cultures.** 4-hydroxybenzoic acid (4-HBA) was added to 10 mL LPU (left) and Delft media (right) in 100 mL shake flasks. The media was inoculated with strain OKYL029 to a starting $OD_{600}$ of 0.05 and incubated for 5 days at 30°C and 220 rpm shaking. Samples for HPLC analysis were taken at the beginning (top) and the end (bottom) of the cultivation. The LPU media also contained 125 mg/L cinnamic acid that was partly converted to 4-HBA. 4-HBA is not degraded in either of the two media during the 5-day cultivation. The small increase in 4-HBA concentration is likely due to evaporation effects throughout the cultivation.

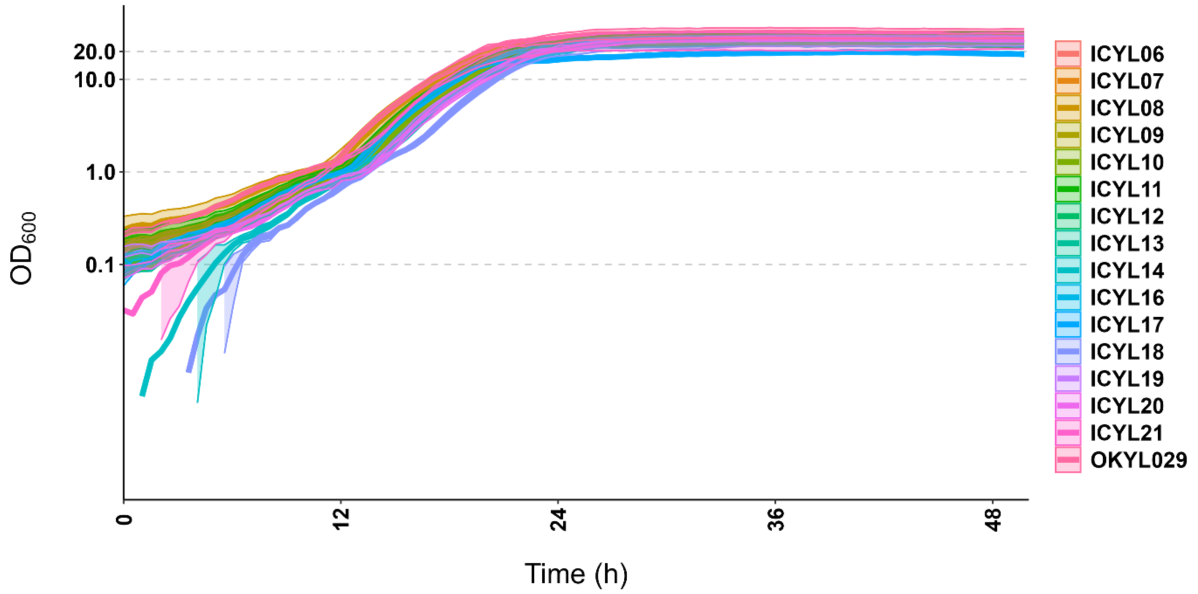


**Figure S4: Growth of the deletion strains in Delft media.** Deletion strains and OKYL029 (as a control) were cultivated in 96-well plates, and pictures to calculate the $OD_{600}$ were taken every 30 min with the Growth Profiler 960. Lines and ribbons represent the mean and standard deviation of four replicates, respectively.

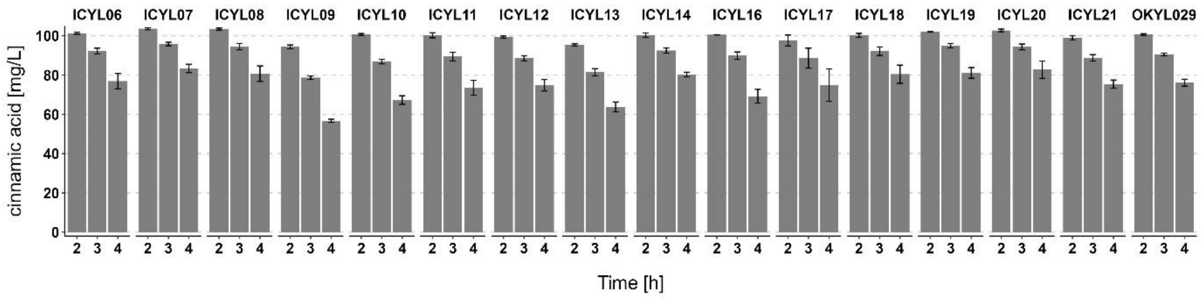


**Figure S5: Cinnamic acid consumption of knock-out strains in Delft media.** Overnight cultures were cultivated in Delft media supplemented with 125 mg/L cinnamic acid and diluted to a starting OD of 0.05 in 10 mL Delft media supplemented with 125 mg/L cinnamic acid in 100 mL shake flasks. HPLC samples were taken at different time points and analyzed. Bars and error bars represent the mean and standard deviation of at least three replicates, respectively.

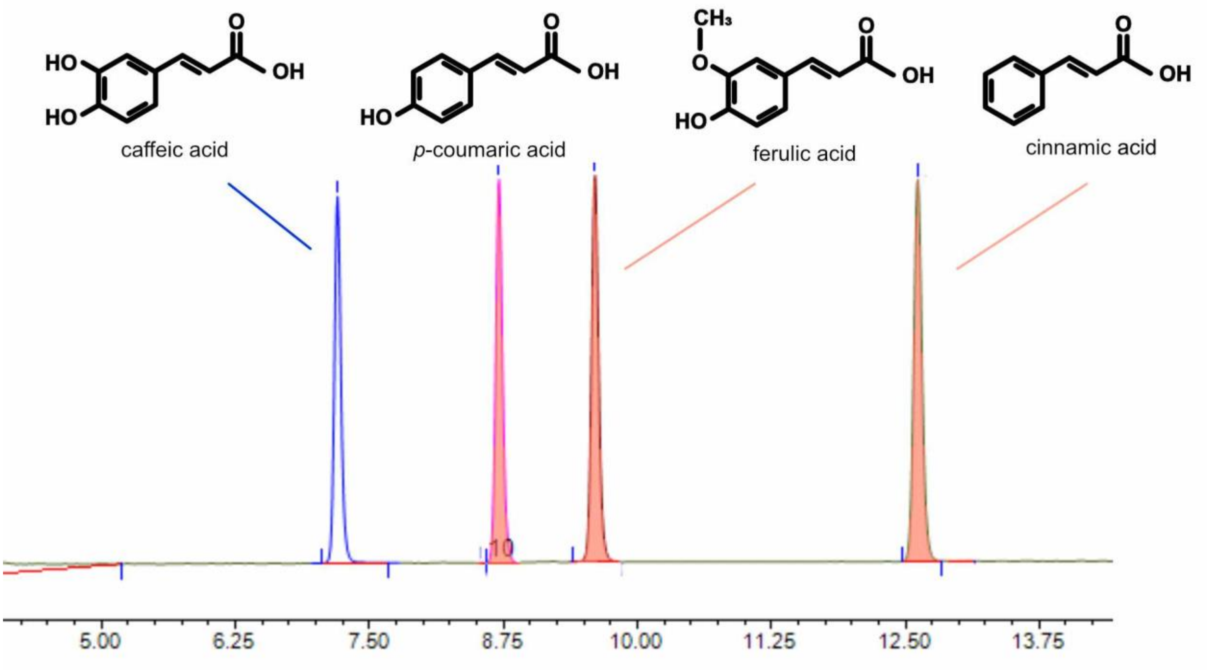


**Figure S6: HPLC chromatogram of the four aromatic acids caffeic, *p*-coumaric, ferulic and cinnamic acid.** The HPLC method used to detect the aromatic acids is described in detail in the method section.

### Supplementary Table

**Table S1: Influence of media composition on *p*-coumaric acid degradation.** Strain OKYL029 was cultured in 10 mL media supplemented with 125 mg/L *p*-coumaric acid in 100 mL shake flasks at 30°C and 220 rpm for 22 h. Cinnamic acid, *p*-coumaric acid and 4-HBA show the average concentration in mg/L ± standard deviation of duplicates. “-“; indicates no detection of the compound in either of the replicates. AS; ammonium sulphate, YaC; yeast extract and casamino acids, VaM; vitamin and trace metal solution.

| **Media base** | **Nitrogen source** | **C/N ratio** | **additional supplements** | **acid added** | **$OD_{600}$** | **cinnamic acid [mg/L]** | ***p*-coumaric acid [mg/L]** | **4-HBA [mg/L]** |
|---|---|---|---|---|---|---|---|---|
| Delft | AS | 3 | | p-coumaric acid | 6.2 ± 0.0 | - | 62.8 ± 2.8 | 38.6 ± 2.9 |
| Delft | AS | 18 | | p-coumaric acid | 11.8 ± 1.0 | - | 54.3 ± 0.4 | 45.1 ± 0.1 |
| Delft | AS | 18 | YaC | p-coumaric acid | 15.9 ± 0.5 | - | 57.0 ± 8.4 | 43.0 ± 2.0 |
| Delft | AS | 116 | | p-coumaric acid | 3.2 ± 0.2 | - | 95.5 ± 0.1 | 16.3 ± 0.2 |
| Delft | urea | 3 | | p-coumaric acid | 6.5 ± 0.1 | - | 85.2 ± 2.2 | 23.5 ± 0.5 |
| Delft | urea | 116 | | p-coumaric acid | 3.0 ± 0.1 | - | 107.3 ± 0.6 | 7.2 ± 0.0 |
| LPU | urea | 3 | | p-coumaric acid | 27.6 ± 0.2 | - | 110 ± 0.8 | 1.9 ± 0.1 |
| LPU | urea | 3 | VaM | p-coumaric acid | 24.6 ± 0.9 | - | 102.9 ± 2.4 | 4.0 ± 0.3 |

| | | | | | | | | |
|---|---|---|---|---|---|---|---|---|
| LPU | urea | 200 | | p-coumaric acid | 18.0 ± 0.1 | - | 93.5 ± 2.9 | 12.8 ± 0.1 |

We compared the different nitrogen sources (ammonium sulphate [AS] and urea), different carbon/nitrogen ratios (C/N ratio), and the addition of either yeast extract and casamino acids (YaC) to Delft media or vitamin and trace metal solution (VaM) to LPU media. In all media, *p*-coumaric acid was degraded and 4-HBA was formed. Suggesting that none of the tested media components alone induces the reaction from *p*-coumaric acid to 4-HBA.

Additionally, the difference in 4-HBA formation in Delft base media can be explained by the different growth ($OD_{600}$) of the different conditions. For example, cultures that grew only to moderate ODs of around 3.0 (delft media with urea at C/N ratio 116) only displayed a 4-HBA production of 7.2 mg/L, while cultures that grew to higher cell densities (Delf media with AS at C/N ratio 18) produced 43 mg/L of 4-HBA.

### Additional supplementary

**<u>Protein BLAST details</u>**

Enzyme: https://www.genome.jp/entry/K00487+1.14.14.91+R02253
From *Arabidopsis thaliana*: https://www.genome.jp/entry/ath:AT2G30490
Protein sequence:

```
>ath:AT2G30490 K00487 trans-cinnamate 4-monooxygenase [EC:1.14.14.91] | (RefSeq) C4H; cinnamate-4-hydroxylase (A)
MDLLLLEKSLIAVFVAVILATVISKLRGKKLKLPPGPIPIPIFGNWLQVGDDLNHRNLVD
YAKKFGDLFLLRMGQRNLVVVSSPDLTKEVLLTQGVEFGSRTRNVVFDIFTGKGQDMVFT
VYGEHWRKMRRIMTVPFFTNKVVQQNREGWEFEAASVVEDVKKNPDSATKGIVLRKRLQL
MMYNNMFRIMFDRRFESEDDPLFLRLKALNGERSRLAQSFEYNYGDFIPILRPFLRGYLK
ICQDVKDRRIALFKKYFVDERKQIASSKPTGSEGLKCAIDHILEAEQKGEINEDNVLYIV
ENINVAAIETTLWSIEWGIAELVNHPEIQSKLRNELDTVLGPGVQVTEPDLHKLPYLQAV
VKETLRLRMAIPLLVPHMNLHDAKLAGYDIPAESKILVNAWWLANNPNSWKKPEEFRPER
FFEEESHVEANGNDFRYVPFGVGRRSCPGIILALPILGITIGRMVQNFELLPPPGQSKVD
TSEKGGQFSLHILNHSIIVMKPRNC
```

| Description | Scientific Name | Max Score | Total Score | Query Cover | E value | Per. Ident | Acc. Len | Accession |
|---|---|---|---|---|---|---|---|---|
| Phenylacetate 2-hydroxylase [Yarrowia lipolytica] | Yarrowia lipolytica | 85.5 | 85.5 | 86% | 1e-17 | 21.66% | 530 | QNQ00524.1 |
| Phenylacetate 2-hydroxylase, putative [Yarrowia lipolytica] | Yarrowia lipolytica | 85.5 | 85.5 | 86% | 1e-17 | 21.66% | 530 | VBB83369.1 |
| YALI0F03663p [Yarrowia lipolytica CLIB122] | Yarrowia lipolytica CLIB122 | 84.3 | 84.3 | 86% | 2e-17 | 21.66% | 530 | XP_504958.1 |
| YALI0E14509p [Yarrowia lipolytica CLIB122] | Yarrowia lipolytica CLIB122 | 69.7 | 69.7 | 40% | 1e-12 | 26.73% | 465 | XP_503945.1 |
| hypothetical protein YALI1_E17508g [Yarrowia lipolytica] | Yarrowia lipolytica | 68.9 | 68.9 | 40% | 1e-12 | 26.24% | 465 | AOW05415.1 |
| cytochrome P450 [Yarrowia lipolytica] | Yarrowia lipolytica | 67.4 | 67.4 | 36% | 6e-12 | 27.72% | 521 | RDW30993.1 |
| YALI0C10054p [Yarrowia lipolytica CLIB122] | Yarrowia lipolytica CLIB122 | 67.0 | 67.0 | 36% | 7e-12 | 27.72% | 521 | XP_501667.1 |
| YALI0A18062p [Yarrowia lipolytica CLIB122] | Yarrowia lipolytica CLIB122 | 65.5 | 65.5 | 35% | 2e-11 | 26.63% | 507 | XP_500188.1 |
| n-alkane- inducible cytochrome P460 [Yarrowia lipolytica] | Yarrowia lipolytica | 58.5 | 58.5 | 33% | 3e-09 | 31.07% | 583 | VBB88954.1 |
| YALI0C12122p [Yarrowia lipolytica CLIB122] | Yarrowia lipolytica CLIB122 | 58.5 | 58.5 | 33% | 3e-09 | 31.07% | 583 | XP_501748.1 |
| hypothetical protein B0I72DRAFT_81777 [Yarrowia lipolytica] | Yarrowia lipolytica | 58.5 | 58.5 | 33% | 4e-09 | 31.07% | 583 | RDW32852.1 |
| hypothetical protein BD777DRAFT_137619 [Yarrowia lipolytica] | Yarrowia lipolytica | 57.8 | 57.8 | 33% | 7e-09 | 30.51% | 578 | RMI95346.1 |
| hypothetical protein YALI1_C17031g [Yarrowia lipolytica] | Yarrowia lipolytica | 57.4 | 57.4 | 33% | 8e-09 | 30.51% | 583 | AOW02734.1 |
| YALIA101S04e11782g1_1 [Yarrowia lipolytica] | Yarrowia lipolytica | 57.4 | 57.4 | 33% | 8e-09 | 30.51% | 583 | SEI34152.1 |
| Cytochrome P450 52A5 [Yarrowia lipolytica] | Yarrowia lipolytica | 57.4 | 57.4 | 33% | 8e-09 | 30.51% | 578 | QNP96247.1 |

The Protein BLAST resulted in the identification of the following 17 target genes, which were deleted to find that YALI1_B28430g is the gene encoding an enzyme with a trans-cinnamate 4-monooxygenase activity similar to the C4H enzyme from *Arabidopsis thaliana*.

| **YALI0** | **YALI1** | **K number** | **Bits** | **E-value** |
|---|---|---|---|---|
| YALI0A15488g | YALI1_A15544g | | 87.8 | 7e-19 |
| YALI0F03663g | YALI1_F05415g | K10437 | 83.4 | 7e-18 |
| YALI0F01320g | YALI1_F02132g | | 83.2 | 2e-17 |
| YALI0B21824g | YALI1_B28430g | | 77.4 | 2e-15 |
| YALI0A20130g | YALI1_A21175g | | 74.3 | 1e-14 |
| YALI0B13838g | YALI1_B18347g | | 72.4 | 4e-14 |
| YALI0E23474g | YALI1_E27649g | | 70.5 | 2e-13 |
| YALI0E14509g | YALI1_E17508g | | 69.7 | 3e-13 |
| YALI0B20702g | YALI1_B27057g | | 69.7 | 4e-13 |
| YALI0C10054g | YALI1_C14106g | | 67.0 | 2e-12 |
| YALI0B01848g | YALI1_B02833g | | 67.0 | 2e-12 |
| YALI0A18062g | YALI1_A18344g | K09831 | 65.5 | 8e-12 |
| YALI0E25982g | YALI1_E30815g | | 62.4 | 6e-11 |
| YALI0C12122g | YALI1_C17031g | | 58.5 | 1e-09 |
| YALI0B13816g | YALI1_B18311g | | 58.2 | 2e-09 |
| YALI0B06248g | YALI1_B08321g | | 51.6 | 2e-07 |
| YALI0B05126g | YALI1_B07094g | | 50.4 | 4e-07 |
